# From *pseudotime* to true dynamics: reconstructing a real-time axis for T cells differentiation

**DOI:** 10.1101/2022.06.09.495431

**Authors:** Avishai Gavish, Benny Chain, Tomer M Salame, Yaron E Antebi, Shir Katz, Shlomit Reich-Zeliger, Nir Friedman

## Abstract

Numerous methods have recently emerged for ordering single cells along developmental trajectories. However, accurate depiction of developmental dynamics can only be achieved after rescaling the trajectory according to the relative time spent at each developmental point. We formulate a model which estimates local cell densities and fluxes, and incorporates cell division and apoptosis rates, to infer the real time dimension of the developmental trajectory. We validate the model using mathematical simulations, and apply it on experimental high dimensional cytometry data obtained from the mouse thymus to construct the true time-profile of the thymocyte developmental process. Our method can easily be implemented in any of the existing tools for trajectory inference.

## Introduction

Technological advancement in measuring expression profiles of multiple variables at single cell resolution has paved the way for creation of new analytical frameworks called trajectory inference. Based on similarity in high-dimensions, dozens of methods exist today that allow sorting cells along the temporal course of biological processes(Cannoodt et al., 2016; Liu et al., 2017; Saelens et al., 2019; Setty et al., 2016; Wolf et al., 2019). Techniques are constantly evolving to address systems of growing complexity, as when cells bifurcate or when development proceeds in a tree-like topology. Recent studies have compared the strengths and weaknesses of these methods to aid users to select the method that best suits their data (Koch and Radtke, 2011; Sawicka et al., 2014; Yui and Rothenberg, 2014). However, trajectory inference, also known as ‘pseudotime’ analysis, can only capture the topology and sequence of the developmental process. In order to capture the time dimension, incorporating the length of time spent by each cell in each developmental stage, additional information on the fluxes within the system (the number of cells entering and leaving the compartments) is required. Several studies attempted to describe true temporal dynamics using some a-priori knowledge about the system (Weinreb et al., 2018), reconstruct the trajectory using experiments to infer some time-points along it (Farbehi et al., 2019; Fischer et al., 2019), or ignored cell division and cell death (Kuritz et al., 2020).

A well-studied system for trajectory inference is the mouse thymus. Bone marrow hematopoietic progenitors migrate to the thymus, where they commit to the T-cell lineage and further mature into functional T lymphocytes(Koch and Radtke, 2011). T-cell development proceeds through a series of discrete phenotypic stages that are characterized by the expression of several important membrane molecules, most notably CD4 and CD8. Thymocytes initially expressing low CD4 and CD8 levels (CD4-CD8-) in the double-negative stage (DN). The DN stage can be subdivided into four distinct phases (DN1 to DN4), each characterized by a different marker expression profile(Germain, 2002; Seo and Taniuchi, 2016; Shah and Zúñiga-Pflücker, 2014). The cells gradually start to express higher levels of both molecules (CD4+CD8+) and transition into the double-positive stage (DP). During the DP stage, cells massively proliferate and undergo two selection processes: positive and negative. In the course of positive selection, only cells that recognize MHC molecules by their T-cells receptors (TCR) survive, while cells that fail to do so die by neglect (Hernandez et al., 2010; Kurd and Robey, 2016; Melichar et al., 2013; Robert et al., 2021; Ross et al., 2014; Szondy et al., 2012; Yamashita et al., 1993). In the course of negative selection, cells that have high affinity to the MHC self-peptide complex undergo apoptosis (Breed et al., 2019; Hernandez et al., 2010). At this point the developmental trajectory bifurcates towards one out of two single-positive stages (SP), where some cells suppress their high CD8 levels while accumulating further CD4 (CD4+CD8-CD3+), or vice versa (CD4-CD8+CD3+)(Figure 1A) (Germain, 2002; Sawicka et al., 2014; Shah and Zúñiga-Pflücker, 2014; Yates, 2014a; Yui and Rothenberg, 2014). During this maturation process, proliferation and apoptosis occur at varying rates, while many more surface markers are upregulated or downregulated upon the differentiating T-cells (le Campion et al., 2002; Chopp et al., 2020; Hernandez et al., 2011; Le et al., 2020; Sawicka et al., 2014; Yates, 2014b). Cell development and selection in the thymus play key roles in shaping the adaptive immune system and maintaining self-tolerance (Goodnow et al., 2005; Griesemer et al., 2010; Hogquist et al., 2005).

**Figure 1:**
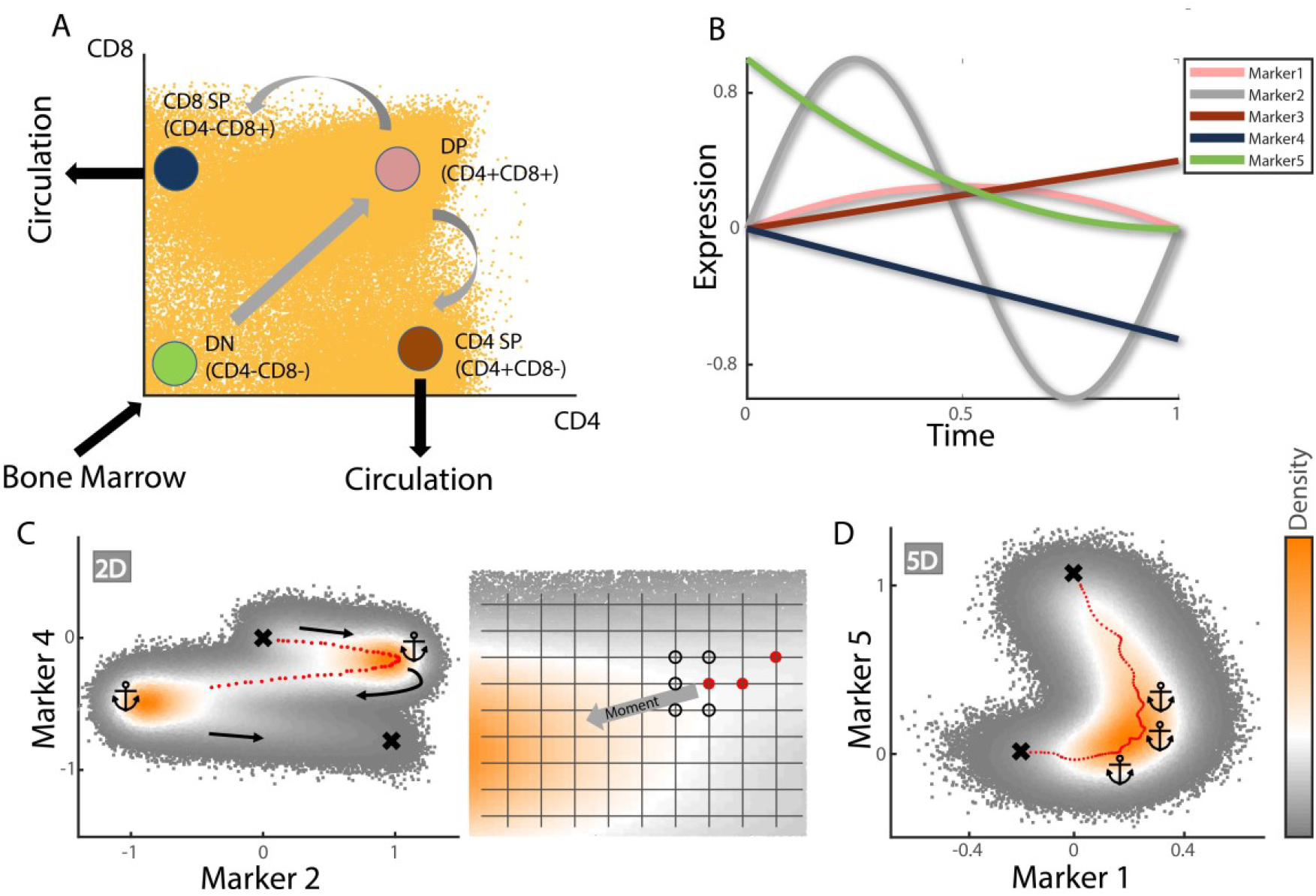
Dimensionality reduction and segmentation. A) Cell differentiation in the thymus. Scatterplot of cells in representative mouse (yellow) depicts a snapshot of the maturation process, with naïve cells entering the thymus in the double-negative state (DN, CD4-CD8-), sequentially differentiating into double-positive cells (DP, CD4+CD8+), and finally acquiring one of the single-positive fates (SP, CD4+/-CD8-/+). Arcsine transformation was used here and for gating purposes (see Methods). B) Five simulated markers. Profiles are shown without adding noise. Each time step (dt=0.01), 100 new cells were added and assigned with t=0. Initial marker values for each cell were chosen from a normal distribution centered at the values plotted at t=0, with a standard deviation 0.1. Noise was also added to the cycle time length of each cell (after which the cell was removed) which was chosen from a normal distribution centered at T=1 with a standard deviation 0.1. Division and apoptosis were not implemented in these simulations, which were run until reaching steady state. C) Trajectory course calculated upon density plot of two markers in B. Starting and ending points are predetermined (upper and lower crosses respectively). Anchors (corresponding symbols) are placed in positions of local maxima. The trajectory (red circles) travels upon the crest connecting path edges and anchors. Panel to the right magnifies the last position obtained and illustrates how new position coordinates are chosen; new coordinates are chosen from nearest neighbors surrounding the previous position on a grid, where progression in the direction of the previous position is forbidden (optional new coordinates are surrounded by black circles). Default choice is for the neighbor in the direction aligned towards the next anchor (moment). Deviation is permitted only if the difference in densities between the default neighbor and some other permitted neighbor exceeds a predetermined value, thereby justifying a detour. See methods for more details. D) Trajectory calculation in five dimensions. The principle is similar to calculation in two dimensions (see text), allowing for trajectory dynamics in any region where at least one of the markers changes. The calculated trajectory is projected here upon a two-dimensional density plot, with anchors and crosses representing landmarks as in C. Note that anchors are placed near, but not in the exact position, of the global maxima in two dimensions (see methods for superfluous anchor removal in higher dimensions).

Here, we use mass cytometry (CyTOF) to measure high dimensional protein expression in thousands of unsynchronized differentiating single-cells simultaneously. Data derived for each cell includes a comprehensive panel of surface marker levels, together with transcription factors and markers for cell-cycle stage and cell-activation. Using ^127^IDU-injected mice (Behbehani et al., 2012), we were able to infer division and apoptosis rates throughout the process.

Similar to previous pseudotime methods, we first formulate an intuitive model for sorting cells along a developmental axis (Kafri et al., 2013; Setty et al., 2016). Our pseudotime ordering method emphasizes the importance of local cell densities, which is a key concept in the consecutive step (however, our real time method can be implemented directly on one of the prevailing methods). Next, by incorporating compartment-specific proliferation and death rates, we capture quantitatively the flow of cells though the developmental process. In this way, we add a true time dimension to the basic developmental sequence and topology captured by previous approaches.

## Results

### Trajectory inference in multidimensional data

The initial step is to order cells or segments of ‘similar’ cells, according to chronological order along the developmental course of events. To illustrate how this method works, we start by simulating an imaginary process where cells enter the system at a constant flux at time *t* = 0, and begin expressing five markers as illustrated (Figure 1B). Random noise was added so that initial values of each marker were chosen from a distribution centered at 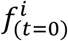 with a standard deviation of 0.1 (*f*^*i*^ is the expression function for each *i* = [1,5] marker). Noise was also added to the entire cycle time length *T* of each cell after which cells were removed, so that *T* was chosen from a random distribution centered at *T* = 1 with a standard deviation equal to 0.1. Assuming no cell division or apoptosis during the process at first, marker values were updated every *dt* = 0.01 time interval. The simulation was run until reaching steady state, in which the total cell number was constant up to some level of fluctuations due to noise. Importantly, at steady state, local cell density as a function of time was also constant (SI Figure S1).

For simplicity, we first calculate the trajectory in two-dimensions, choosing the markers illustrated in Figure 1C. Marker expression space is divided into a *m*^*i*^ · *m*^*j*^ grid, with *D*^*ij*^ denoting cell density at each grid position. Our working assumption is that the trajectory must pass through points of local maxima in cell density (anchors in Figure 1C). We also assume that the initial and final positions of the trajectory are known approximately and are manually determined (e.g. the assumed location of the starting and ending cells). The trajectory, denoted as *l*, then progresses between points on the grid from one anchor to the next, traveling upon the density crest. The default trajectory is in the direction of the next anchor (‘moment’), while a detour is only permitted when the difference in densities between the grid point aligned with the moment and that at some other neighboring point exceeds some predetermined value (See SI section 2 for more details).

Computing trajectory coordinates *l*^*ij*^ effectively reduces dimensionality to one. Each cell is now associated with a single point upon *l* to which it is closest based on Euclidean distance. Assuming the trajectory is composed of *k* evenly spaced points, each point can generaly affiliate many cells; the *k* cell-segments are ordered according to the chronological course of the systems’ evolution. Inside a given segment, there is no internal ordering of cells. Thus, the number of sections (or initial grid density) and the extent of trajectory curvature, dictate the resolution by which cells are sorted.

Trajectory inference in this method is closely dependent on dynamics in marker expression. E.g., if the two markers in Figure 1C happen to be stationary at some point, density in two-dimensional space around that point will be relatively high. Extending the method to higher dimensions by adding more markers is therefore beneficial, as local kinetics in expression of even a single marker is able to unfold regions where seemingly no dynamics occur. Following a similar line as in two-dimensions, the trajectory can be calculated in higher dimensions by dividing each dimension into a grid and passing from a given initial position through regions of local maxima. Notably, removal of superfluous local maxima points might be necessary after expanding the number of dimensions (SI section 2). Figure 1D illustrates the trajectory that was calculated in 5-dimensions (using all 5 markers) projected upon a two-dimensional density plot.

### From pseudotime to real-time: theory and simulation

The method described above segments similar cells according to Euclidian proximity in high dimensions with respect to the trajectory. To illustrate the principle by which real-time can be calculated, it is useful to first focus on fluxes and dynamics within a single segment of cells (Figure 2A). Within every time interval, a fraction of cells mature to a point where they exit the segment and enter the neighboring one ahead. Similarly, some other fraction of cells enters the segment by exiting the preceding segment. For a segment with index *i*, these density fluxes of entering and exiting cells can be written respectively as *f*_*i*−1_ · *ν*_*i*−1_ and *f*_*i*_ · *ν*_*i*_, where *f* denotes cell density (the number of cells in a segment divided by the total number of cells), and *ν* denotes the progression rate along the trajectory *l*. Other sources for cell turnover within a segment are cell division and apoptosis, whose rates are denoted as *γ*_1_ and *γ*_2_respectively. Every time interval, 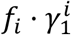 cells are added to the segment, while 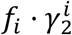 cells are removed. Thus, the local density within a segment, which in steady state must remain constant over time, is ultimately defined by the different fluxes of entering and exiting cells, and the local proliferation and apoptosis rates. A high local density, in the presence of a low division-to-apoptosis rate ratio, is indicative of a prolonged dwelling-time of cells in the segment, and vice versa. This conservation principle is presented compactly in the equation shown Figure 2B. A closed-form expression in the case when *γ*_1_ and *γ*_2_ are constant in space is highlighted below the equation (rectangle), where *t* is the cumulative average time spent up tp point *l* on the trajectory, and represents the avarge developmental time of cells in that segment (More derivatives of this equation are given in SI section 3).

**Figure 2:**
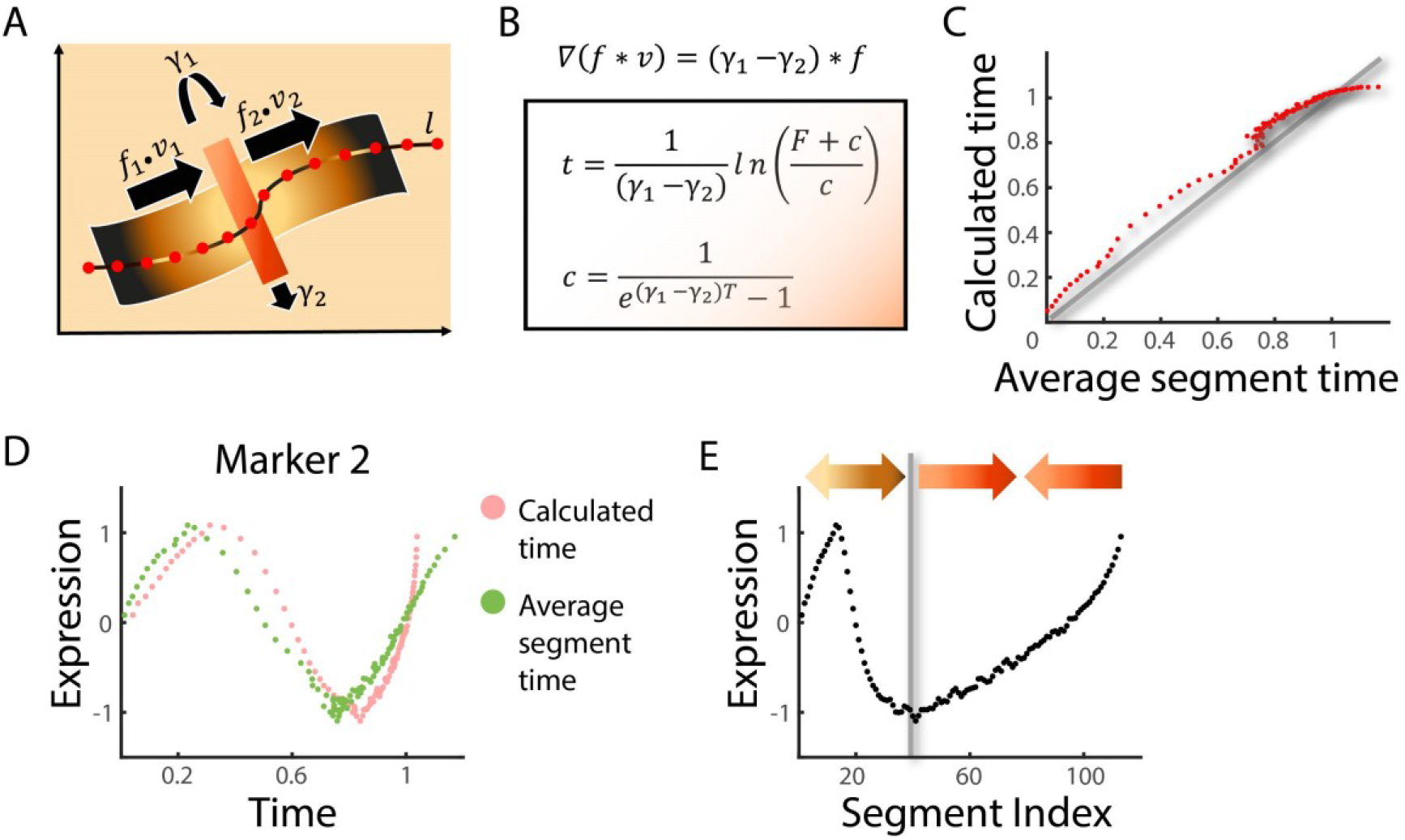
Time calculation. A+B) The time equation. Cartoon outlining dynamics within a single segment is shown in A. Red circles represent trajectory points in some imaginary data, with the red stripe magnifying a given segment. The local density in each segment is determined by the flux of entering cells *f*_1_•*ν*_1_ (*f* and *ν* represent local cell density and velocity, respectively), flux of cells exiting the segment *f*_2_•*ν*_2_, and cell division and apoptosis rates (*γ*_1_ and *γ*_2_, respectively). High local density can be a manifestation of lengthy dwell-time of cells in a segment, or of a high division to apoptosis rate ratio. Similarly, low local density can be a result of rapid progression through the segment, or of a high apoptosis to division ratio. B gives the time equation in its general form with ∇ denoting the differential operator. The formulation for the time as function of location along the trajectory, *t*(*l*), is given after solving the time equation assuming constant *γ*_1_ − *γ*_2_ > 0, with *F* denoting the cumulative density up to location *l* = *l*’, and *T* denoting the time reached when cells arrive at the trajectory end. See methods for more derivations including numeric formulation for when division is not constant. C) Validation of formulation accuracy. Simulations were performed as in Fig 1B, this time incorporating constant division and apoptosis rates (*γ*_1_=0.09, *γ*_2_=0.03). The ‘Average segment time’ directly averages simulation times of cells in each segment and represents real time. ‘Calculated time’ represents time in each segment using the formulation in B. The deviation towards the end is due to an increasing number of cells exiting the cycle, with the few remaining cells adding negligibly to the cumulative density. Correlation coefficient reaches 0.99. D) Average values of Marker2 (Fig 1B), using real segment average time (green) and calculated times (pink). E) Average marker times as function of segment index (*pseudo-time*) distorts the expression profile. Arrows indicate re-scaling applied by the time equation.

To test the theory formulated above, we simulated the dynamics in Figure 1B, this time adding proliferation and apoptosis. Figure 2C compares the ‘real’ dwell time, which was calculated by averaging the known simulated times of all cells within each segment, to the time calculated using the equation given in Figure 2B. The simulation and prediction agree to within a threshold determined by noise (Figure 2C), and by a deviation as t approaches 1 that can be explained by the open-ended model, allowing cells to leave the system, rather than dying. The trajectory of marker expression, plotted as a function of calculated or simulated times are also in good agreement (Figure 2D), but differ substantially when plotted as a function of pseudotime, which does not incorporate division and death methods (Figure 2E).

### Experimental determination of proliferation and apoptosis rates

To measure cell proliferation in-vivo, we injected mice with 5-Iodo-2′-deoxyuridine (^127^IDU), a thymidine analog incorporated to DNA during S-phase. Accordingly, only proliferating cells incorporate IDU and integrate it into the DNA. Upon further division, half of the initially incorporated IDU is passed on to each daughter cell. Following injection, we harvested the mouse thymus after 1, 4, 6 and 12 hours (3 mice in each group) and performed CyTOF analysis (Figure 3A). IDU levels are readily detected as elemental Iodine by CyTOF, with no additional experimental requirements.

**Figure 3.**
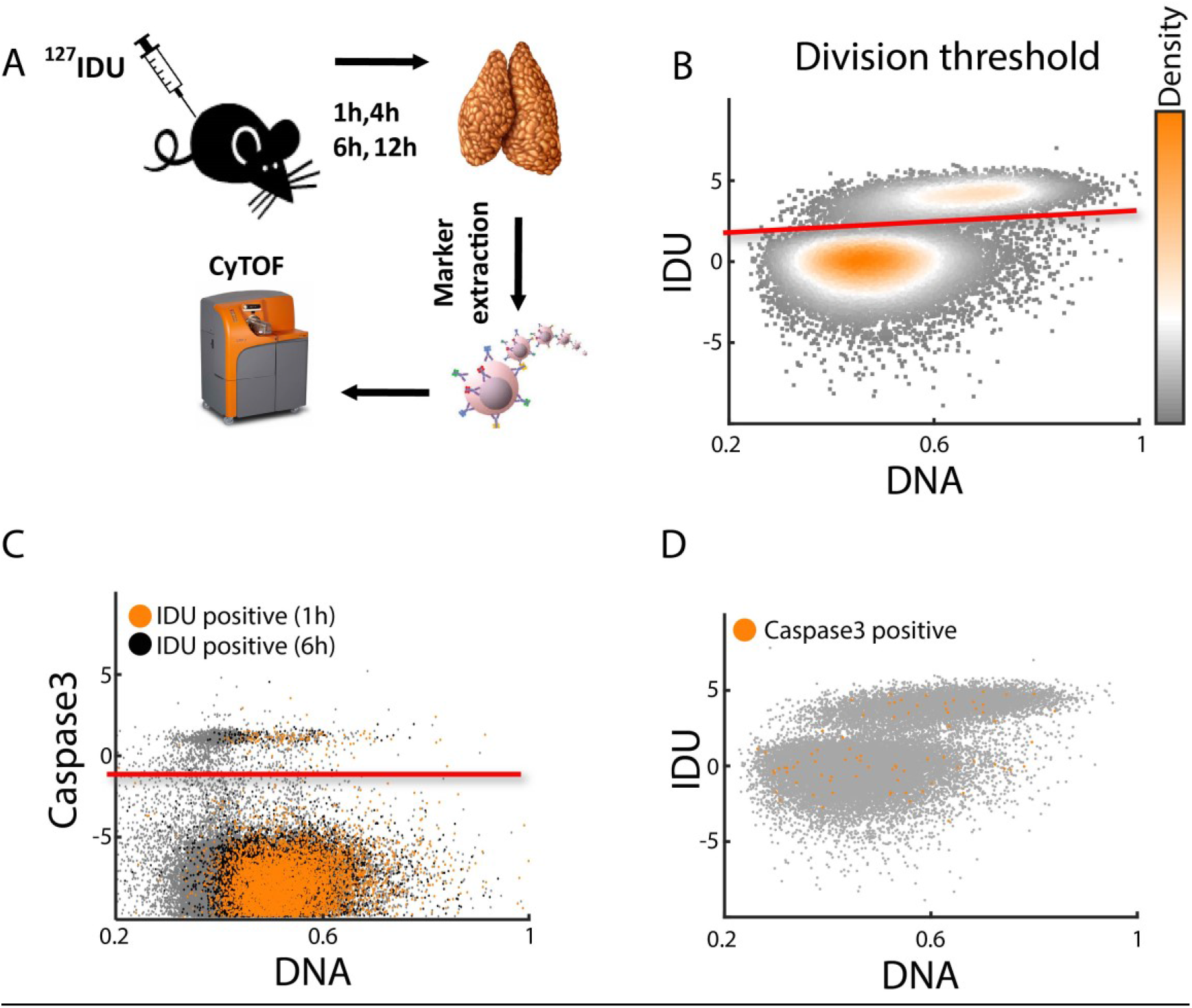
Determining division and apoptosis rates A) Scheme describing the IDU injection process. B,C) Threshold for dividing and apoptotic cells. The logarithm of markers used to estimate the thresholds for division and apoptosis (IDU and Caspase3, respectively) are plotted against normalizes DNA. Panel B shows data from a mouse 1 hour post IDU injection, and panel C shows data from two mice 1 and 6 hours post IDU injection. Cells above the red line in B were considered as dividing cells within 1 hour. Cells above red line in C, that were also positive to IDU, were considered as cells that will undergo apoptosis within 1 or 6 hours. *γ*_1_ and *γ*_2_ in each segment for this mouse was calculated as percentage of IDU or IDU-Caspase3 positive cells in the segment divided by the corresponding *δt* in day units. D) Caspase3-positive cells in a mouse 1 hour post IDU injection are roughly evenly distributed between the two populations of IDU positive and negative cells (SI Section 4).

Plotting IDU expression as a function of DNA levels, allows identification of two cell populations – dividing and non-dividing cells, with and without IDU uptake respectively (Figure 3B). Gating the dividing (IDU positive) cells, selects the cells that have divided in the time interval that lapsed between IDU injection and thymus harvesting (we denote this time interval as *δt*). We assume that the number of cells dividing more than once within *δt* (max 12h) is negligible. Thus, the proliferation parameter in each segment 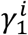 can be calculated by dividing the proportion of IDU-positive cells within the segment by *δt*.

Our chosen apoptosis marker was Caspase3, a 32 kDa cysteine protease that is activated during the early stages of programmed cell death (Tyas et al., 2000). Caspase3-positive cells could be observed as a scatter of cells above the remaining non-apoptotic cells (Figure 3C). Since calculating the apoptosis parameter necessitates dividing by a time interval within which apoptosis will occur, we chose only cells that were positive for both Caspase3 and IDU. To check whether these cells indeed reliably represent the entire population destined for apoptosis within *δt*, and not some unique sub-population, we verified that cells considered Caspase3-positive are evenly distributed throughout the IDU-DNA plane, and are not skewed into some specific region (Figure 3D).

Hence, the gated cells positive to both Caspase3 and IDU were assumed to undergo apoptosis within the same *δt* used for assessing the division rate. Similar to 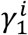, the apoptosis parameter in each segment was calculated by dividing the proportion of IDU-Caspase3-positive cells within the segment by *δt*. Assessing the proliferation and apoptosis parameters should be relatively robust to the time of IDU injection (which is arbitrary). Indeed, the relative population of IDU-positive cells roughly increased linearly with *δt*, resulting in comparable similar proliferation and apoptosis parameters irrespective of injection time (Figure 3C and SI Figure S3).

It is well established that in a short time interval during the positive selection process that occurs in the DP phase, approximately 90% of the thymocytes undergo death by neglect (Sawicka et al., 2014; Stritesky et al., 2013; Szondy et al., 2012). The direct measurement of death-by-neglect using established apoptosis markers such as Caspase3 is nevertheless challenging, since dying cells are rapidly removed by scavenger macrophages, leading to significant under-estimates of the number of dying cells. We verified this extensive apoptosis incident by an in-vivo experiment in which we adoptively-transferred DN thymocytes from CD45.1 (B6SJL) mice into the thymus of CD45.2 (C57Bl) mice, and followed their development (Figure 5A).The observed reduction in CD45.1 cell-number reduction between 21 and 28 days supported the hypothesis that about 90% of cells are eliminated (Figure 5B and SI Figure S3), Based on the literature, and these additional experiments, we therefore modified the parameter controlling the rate of cell death *γ*_2_ in the DP phase so as to model the loss of 90% of the thymocytes during this stage (SI section 3).

**Figure 4:**
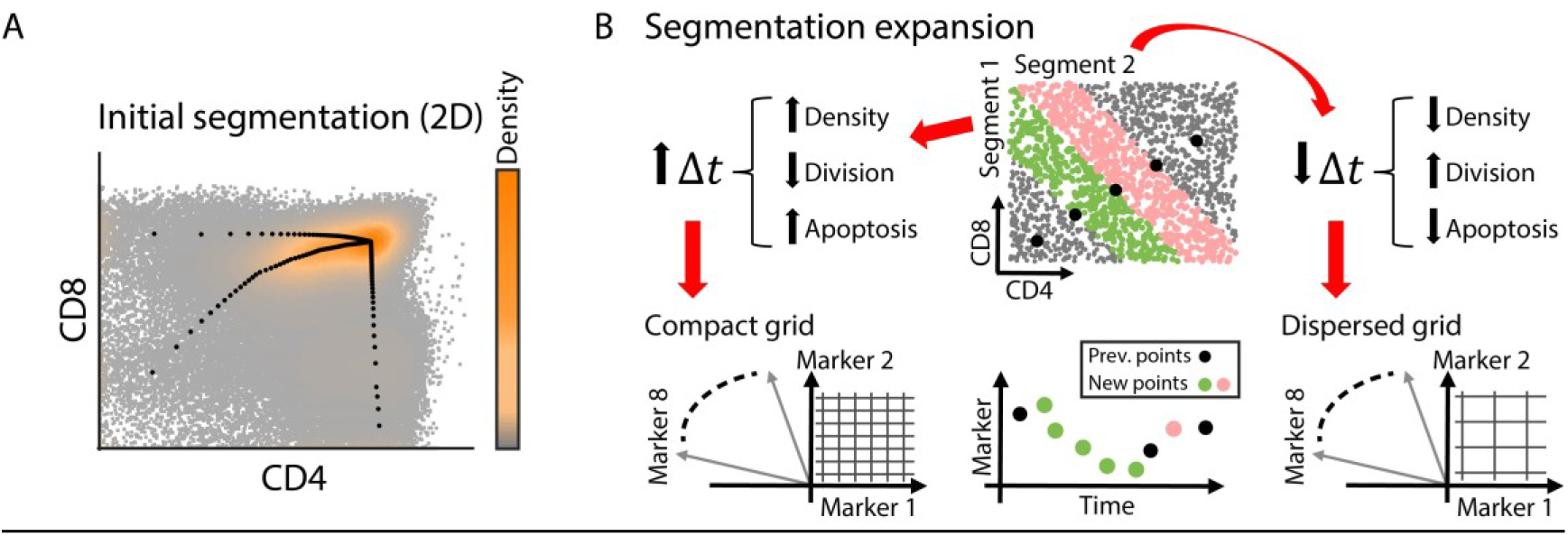
A two-step segmentation process. A+B) Since one-step segmentation in higher dimensions can lead to significant dispersal of cells along the projected trajectory in CD4/CD8 space (e.g. assigning DN cells to points far along the trajectory, SI figure S4), segmentation was first performed in 2D. Panel in A shows the trajectory upon the density plot in CD4/CD8 space, leading from DN to DP, and splitting towards SP states. The time equation was solved separately for the DN+DP phase and each SP phase. Sequentially, each segment was sub-segmented using more markers (8 dimensions) as illustrated in B. A zoomed portion of the trajectory calculated in A is shown, with cells between two pairs of points colored differently (green and pink). The grid size for sub-segmentation was proportionally chosen in respect to the relative time spent in the original segment (Δ*t*), potentially resulting in addition of more or less time points.

**Figure 5:**
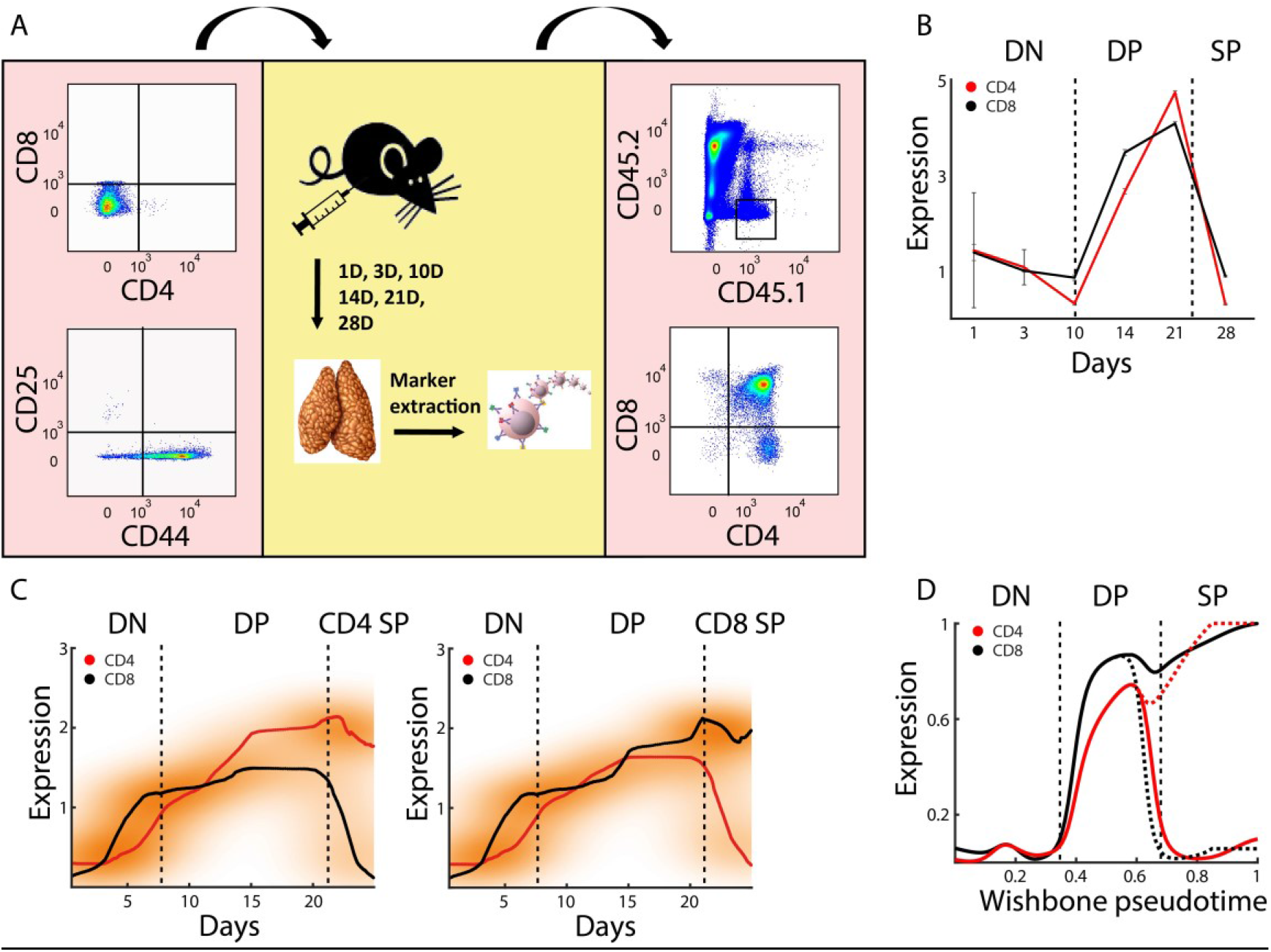
Real-time axis validation A) *In-vivo* validation was performed by intrathymic, adoptive transfer of DN1-3 (CD4^-^CD8^-^CD44^+/-^CD25^+/-^) cells from CD45.1 mice (B6SJL) into CD45.2 (C57BL) mice (left and center panel). Thymi were harvested 1, 3, 10, 14, 21 and 28 days post transfer and stained with antibodies against CD45.1, CD45.2, CD8, CD4, and other developmental markers (center). Further analysis was performed by gating the CD45.1+/CD45.2-cells. Each time point presents Thymi from two mice, and the experiment was repeated twice. B) CD8 (Black) and CD4 (Red) expression levels were calculated tracking the CD45.1+/ CD45.2-gated cells. The expression levels for CD4 and CD8 were separately normalized to the median fluorescence of all the cells from all the time points. Black broken vertical lines separate here (and in later figures) between the DN, DP and SP phases as indicated on top. C) CD4 and CD8 profiles as constructed by our method for real-time axis inference averaged across 12 mice. D) Wishbone pseud-temporal ordering captures the major stages in T-cell development but scales differently than the real-time axis.

### Imputation of a real time dimension in the thymus developmental trajectories

CD4 and CD8 are the two most pivotal markers that define the final outcome as well as key transitional phases along a thymocyte developemental trajectory. Trajectory calculation in high dimensions and cell-affiliation to trajectory points according to Euclidian distance can result in cell dispersion across CD4/CD8 space, so that for instance some cells expressing low CD4 and CD8 (and therefore considered to be in DN phase) are assigned to relatively distant points along *l* (SI Figure 4). This results from the fact that a simple Euclidian calculation gives all markers a similar weight in defining cell proximity to trajectory points. A priori knowledge of CD4 and CD8 importance in specifying differentiation states can therefor help refine the segmentation process.

We thus performed trajectory calculation and cell assignment in two steps. At first we calculated the trajectory in CD4/CD8 plane and assigned cells to trajectory points using these two markers alone (Figure 4A). The bifurcation point was identified manually, along with the initial and final trajectory points. The estimated dwelling time of cells in each segment was calculated as described above. We next performed sub-segmentation by considering more markers. For each given segment, aside from CD4 and CD8, we chose another six markers with the highest local variability in the segment, which thus have the potential of contributing the most to understanding the systems’ dynamics. In segments where cells dwelled for a relatively long period according to the calculation in 2-dimensions (high Δ*t*), we wanted a higher resolution, and therefor divided the segment into more points by choosing a compact grid. On the other hand, in regions where cells spent less time according to the calculation in 2-dimensions (low Δ*t*), we allowed a sparser grid and hence less points were added within the segment (Figure 4B).

Dividing the segmentation process into two steps enforced the distribution of cells according to the CD4/CD8 expression. Sub-segmentation in the second step allowed ‘unfolding’ of regions where additional transitions could be discovered by addition of more markers.

Construction of the time axis allows comparison of different marker profiles along the two SP branches. CD4 and CD8 expression profiles follow the expected trend in which both markers rise up to the bifurcation point (end of DP phase), after which the counter marker starts to decrease along the respective branch (Figure 5C). To compare our expression profiles to those obtained using pseudotime, we used the well established algorithm Wishbone, originally used to infer developmental trajectories for maturing thymocyte(Setty et al., 2016) (Figure 5D). The two approaches were successful in detecting the major developmental states, but the relevant time spent in each state was apparently different. While the Wishbone trajectory suggested that cells spent ∼30% of the time in the DN state and ∼20% of the time in the DP state, our real time trajectory indicates that cells spent less than 20% in the DN state, and about 60% of the time in the DP state. One limitation of our segmentation approach is in detecting relatively rare cells whose population size is much lower than the number of cells in the segment to which they belong. For example, the DN1 (CD4-/CD8-/CD3-/CD25-/CD44+) population size is considered to be about 1% of all DN cells (and ∼0.03%-0.05% (Laurent et al., 2004)), while the initial segments in the trajectory are more than an order of magnitude larger in terms of cell numbers. Indeed, the initial segments in our model seem to capture the dynamics as of the DN3 phase in which cell density starts to increase rapidly, which results in an effective shortening of the DN phase. This is seen in Figure S5 and S6 where CD25 and CD3 levels are relatively high, consistent with the DN3 phase (CD4-/CD8-/CD3+/CD25+/CD44-). To validate the predicted dwell-times in the different developmental phases predicted by the model we sought to determine the timeline of differentiation experimentally. To this end, we used the adoptive-transfer model outlined above, detecting T-cells 1,3,10,14,21 and 28 days post transfer (Figure 5A). We observed that CD45.1 T cells remained in the DN state for approximately 7 days, after which they began to express CD4 and CD8 in parallel, thereby entering the DP state. About 25 days post transfer we could not detect ∼90% of cells. We speculate that the reduction in cell number was since cells either underwent apoptosis (via positive or negative selection), or matured to CD4 or CD8 SP cells and left the thymus (Breed et al., 2019; Sawicka et al., 2014; Stritesky et al., 2013). The inferred dwell-times therefore correspond well to the experimental observations, and fit the data better than the standard pseudotime projections.

The full profiles of all markers we used can be found in SI Figure S6.

### A real-time trajectory recognizes dynamics overseen using pseudotime

We examined the inferred dynamics of expression of all the markers, and compared them to the Wishbone trajectories. The general trends were similar for most markers (SI Figure S6). However, some specific differences emerged. CD69, a marker of TCR signaling(Ross et al., 2014; Yamashita et al., 1993) exhibited complex dynamics during the DP stage, with a high expression peak around day ten, which declined sharply and then rose again between days 15-20 (Figure 6A). The second peak was was less obvious in the Wishbone trajectory (Figure 6A, right panel).

**Figure 6:**
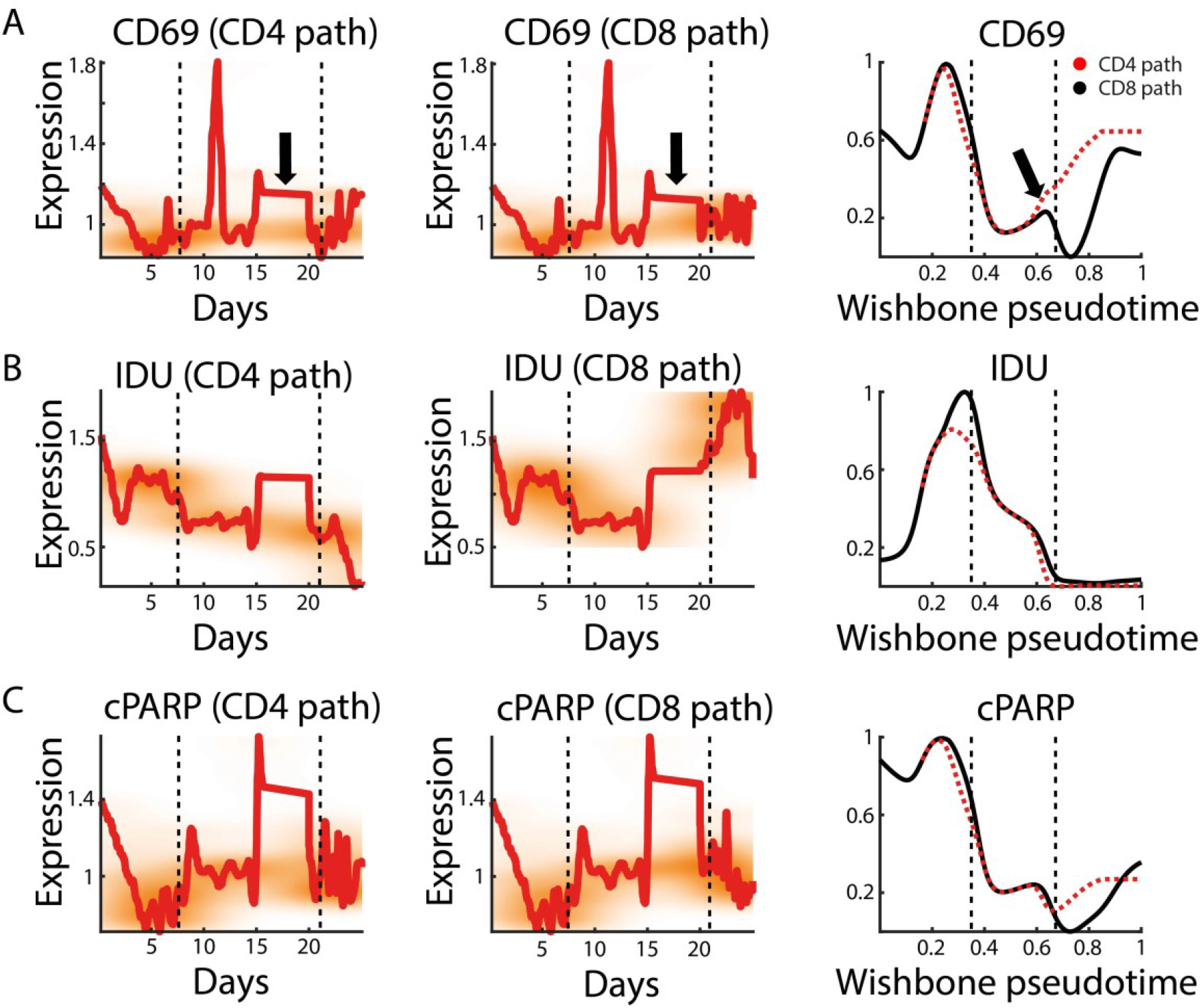
The real-time axis is able to unfold regions of biological significance A,B,C) CD69, IDU and cPARP real time trajectory profile along the CD4 SP and CD8 SP paths (left and center panels respectively), and along Wishbone pseudotime trajectory (right). Black broken vertical lines separate between DN, DP and SP phases as in figure 5. Black arrows in A indicate the second elevation in CD69 expression which is spread upon several days along the real-time axis as opposed to the relative condensed portion it occupies in the Wishbone trajectory.

A similar but delayed pattern is seen in expression of cleaved-PARP1 (cPARP), which is produced during both apoptosis and necrosis(Elmore, 2007) (Figure 6C). Proliferation, identified by IDU incorporation, also exhibits a bi-wave profile, where cells divide more extensively around day 5 during the third phase of the DN state (DN3)(Laurent et al., 2004; Paiva et al., 2021), and then again at the DP state(Ciofani and Zúñiga-Pflücker, 2007; Li et al., 2021) between days 15 and 20 (Figure 6B).

CD25 and CD44 are two markers which are known to have higher expression levels during the DN phase and decrease during the DP phase (SI Figure S5). Notably, the profiles of CD25 and CD44 were inferred primarily via dynamics in expression of other markers who defined the trajectory route, as these two markers did not happen to participate in sub-segmentation of any of the segments. Interestingly, while CD25 seems to decrease towards the DP phase and remain relatively low in the SP phase along both branches, CD44 decreases during the DP phase along the CD8-SP branch but rises towards the end of the CD4-SP branch.

The method described in this paper not only infers the experimentally observed dynamics of the overall developmental trajectory, but identifies and highlights some more complex dynamic features of individual molecules which are compatible with the known complex biology of thymocyte differentiation.

## Discussion

Trajectory-inference has gained wide popularity as a tool for analyzing single-cell omics data. Although many ordering approaches have evolved during the last few years, most methods are comprised of some sort of dimensionality reduction followed by trajectory inference based on expression profile similarities, and ultimately allocate cells along a continuous pseudo-temporal axis(van den Berge et al., 2020; Cannoodt et al., 2016).

Our study builds on this work by estimating cell fluxes along the developmental trajectory, and hence rescaling the pseudotime axis to more closely reflect the real dynamics of thymic differentiation.

We initially formulate a method for pseudo-temporal ordering based upon expression-densities, which is conceptually similar to existing methods but highlights the importance of local cell densities as preparation for the next step. While this method is less sensitive in detecting relatively rare cell populations that can be diluted within segments (such as the DN1 and DN2 cell populations), it is less sensitive to initial condition selection (e.g. as seen in cases where an initial single cell is selected). We further incorporate apoptosis and division, and assuming steady-state, infer the real time dimension.

Developing T-cells in the mouse thymus are an ideal system for trajectory-inference, since cells are known to continuously travel across developmental stages (characterized by different markers) in a well defined period of time. Implementing our method, we were able to depict the observed course of T-cell differentiation and extract temporal profiles for all markers that we measured. Importantly, regions of the developmental landscape that were compacted by traditional pseudotime inference could be expanded by incorporating additional markers revealing events of biological significance. This temporal reconstruction of the differentiation pathway can readily be applied to other complex developmental systems, and the trajectories can be further refined as higher dimensional data sets become available.

## Material and Methods

### Mouse thymus data and mass cytometry

Female C57BL/6 and B6SJL mice were obtained from Harlan Laboratories. All mice were housed at the Weizmann Institute in compliance with national and international regulations. Thymocytes were isolated from the thymus of 6-week-old C57Bl mice. Cells were stained with metal-conjugated antibodies according to manufacturer’s protocol (Supplementary Table 1). Briefly around 200k cells were stained with cell-ID TM Cisplatin (Fluidigm) (5min RT). Next cells were stained with surface antibodies (30 min RT), and fixed with 1.6% PFA (10 min RT). After permeabilization with 100% ice-cold Methanol (15 min, 4C), the cells were stained with intracellular antibodies (30 min, RT). Finally the cells were labeled with Iridium DNA intercalator for DNA content and analyzed by CyTOF mass cytometry using CyTOF2. Data was normalized using bead normalized with bead standards.

We collected data for 12 independent thymuses from Black6 mice.

### IDU injection

IDU (*5-Iodo-2’-deoxyuridine*) was purchased from Sigma-Aldrich (ref. I7125). All mice were injected in their peritoneum with a solution of 0.4mg of IDU diluted in 200*μl* of sterile PBS, adjusted to a pH of 8.5–9. Thymocytes were isolated 1, 3, 6 and 12h post injection. Cells were stained with metal-conjugated antibodies according to manufacturer’s protocol (Supplementary Table 1). Briefly, around 200k cells were stained with cell-ID TM Cisplatin (Fluidigm) (5 min RT). Next cells were stained with surface antibodies (30 min RT), and fixed with 1.6% PFA (10 min RT). After permeabilization with 100% ice-cold Methanol (15 min, 4 °C), the cells were stained with intracellular antibodies (30 min, RT). Finally, the cells were labeled with Iridium DNA intercalator for DNA content and analyzed by CyTOF mass cytometry using CyTOF2. Data were normalized using bead normalized with bead standard.

### CD45.1 Thymocyte adoptive transfer

Thymucytes were purified from B6SJL (CD45.1) mice. The cells were stained with CD4, CD8, CD44 and CD25 and sorted to the CD4-CD8 double negative population. 10,000 cells in 50ml of the DN population were adoptively transferred directly to the thymus of each C57BL/Hj (CD45.2) mice. Thymocytes were isolated 1, 3, 10, 14, 21 and 28 days post adoptive transfer. Cells were stained with various antibodies and analyzed by Flow cytometer.

## Code and Data availability

Any relevant code or data are available upon reasonable request from the corresponding author. Future versions of the manuscript will also include the code and interface for downstream analysis.

## Acknowledgments

We thank Naama Barkai (Weizmann Institute of Science, Israel), for her help in theoretical design of the model. We thank Manu Setty (Fred Hutch, USA) for his helpful advice on reconstructing the Wishbone trajectory. We thank Erez Greenstein (Weizmann Institute of Science, Israel) for his helpful feedback.

