## Supplementary for "From *pseudotime* to true dynamics: reconstructing a real-time axis for T cells differentiation"

**Table of contents**

Section 1: Marker simulations

Section 2: Trajectory calculation

Section 3: Time calculation

Section 4: Division and apoptosis rates assessment

Section 5: A two-step segmentation process

Section 6: Profiles of all markers

Section 7: Data processing and parameters

### Section 1: Marker simulations

This section refers to simulations shown in Figure 1B we used for initial validation of the model in Figure 2 in the main text. A constant flux of 100 cells per time unit (*dt*=0.01) were added to the system. At simulation time *t*=0, each cell began expressing 5 markers represented by the following functions:

$$Marker1 = -t^2 + t$$

$$Marker2 = \sin(2\pi t)$$

$$Marker3 = 0.4 \cdot t$$

$$Marker4 = -0.65 \cdot t$$

$$Marker5 = t^2 - 2t + 1$$

Noise was initially added to the simulations in two forms. First, at *t*=0, initial marker values were chosen randomly form a normal distribution centered at $Marker_{t=0}^{i}$ (namely the original value of each marker with index $i$ at *t*=0), with a standard deviation of 0.1. Second, the time after which cells were removed from the system, $t = T$, was chosen randomly for each cell from a normal distribution centered at $T = 1$ with a standard deviation equal to 0.1.

In Figure 1 in the main text, we ran the simulations as described, without cells dividing or being removed during the process via apoptosis. In Figure 2, we incorporated a division rate of 9% for all cells each time interval, notated as $\gamma_1 = 0.09/dt$, and an apoptosis rate of 3% for all cells each time interval, notated as $\gamma_2 = 0.03/dt$. Each time step, a cell was designated to division or apoptosis according to a random number generated from a binomial distribution specified by the probability of success (0.09 for division and 0.03 for apoptosis). Thus, some variations in division and apoptosis rates incorporated a further form of stochasticity into the system. Simulations were ran until reaching steady state, where the total number of cells in the system was constant up to some fluctuations due to noise (Figure S1A). Importantly, at steady state local cell density was also constant over time, allowing application of the conservation law at the level of each local segment we later used for calculating time (Figure S1B).

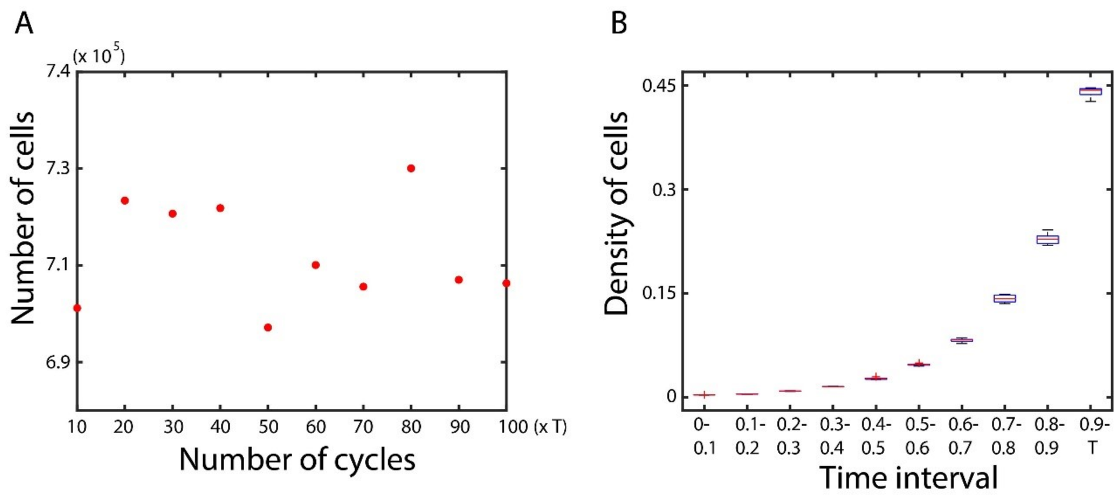


Figure S1: Simulations at steady sate. (A) At steady state the total number of cells in the system is constant up to some level of fluctuations due to noise. Horizontal axis represents time in units of cycle length T. (B) Box plot showing the median density of cells in different intervals. Cells were sampled at steady state every 10 cycles (10 times altogether) at 10 time intervals. The bottom and top edges of each box indicate the 25th and 75th percentiles, respectively. The number of cells grows exponentially with time interval, since the division to apoptosis rate ratio was larger than 1.

### Section 2: Trajectory calculation

Our goal is to convey a trajectory composed of discrete points according to chronological developmental order that also passes in proximity to most of the data (for later accurate assignment of each cell to a trajectory point).

It is useful to first examine trajectory inference in two dimensions where local densities can be easily depicted (Figure 1C). Data is divided into a $m^i \cdot m^j$ grid and cell density is calculated at each grid point. Points of local maxima on the grid, indicated by anchors in the figure, are regarded as landmarks through which the trajectory path must travel. For maximal overlap of the trajectory with the developmental contour, the path connecting the anchors primarily travels upon the density crest connecting maxima points. Given a trajectory point, to determine the position of the next point we compared cell density in grid locations of all nearest neighbors. The preference was choosing the position aligned with the direction of the next anchor ('moment'), while positions within $\pm 90^{\circ}$ in relation to the previous point were forbidden to prevent formation of loops (Figure 1C magnified panel). We did allow deviation from progression towards the next anchor when the density crest curved sharply, that is when the difference between cell-density at the point aligned with the moment and the density at some other permitted grid point exceeded some predetermined value justifying a detour (see Figure S2 and Section 7 below).

To calculate anchor positions, we first used a code written by Stephen M. Anthony, available at:

https://www.mathworks.com/matlabcentral/mlc-downloads/downloads/submissions/45338/versions/2/previews/findMaxima.m/index.html

After obtaining local maxima positions, there are two further steps required before determining the final anchor locations:

1) Superfluous anchor removal. Aside from data smoothing which might prevent selection of potential erroneous anchors that do not represent real regions of local maxima (but rather local fluctuations), calculation in high dimensions can also result in many unwanted points. With addition of more markers, the density grid becomes sparser and points on the grid are more susceptible to having higher density compared to neighboring points (and thereby being selected). We removed superfluous anchors by repeating the maxima search 10 times, each after adding 5% random noise, and choosing points that did not change location in at least 7 runs. While the superfluous points were not robust to this small amount of noise, the true positions of local maxima persisted in most runs.

2) Notably, the points of local maxima are not ordered. Hence, we first directly connected all possible anchor permutations, divided each alternative into the same number of evenly spaced points, and summed the densities at all points. We tried several options for even spacing, to avoid accidental error resulting from differences in path lengths connecting the points (where more local maxima points happen to overlap with a specific spacing of an incorrect permutation). We chose the permutation that consistently summed up to the maximal density. We double-checked our choice by affiliating cells to closest points upon each trajectory and

summing all distances. Indeed, the chosen trajectory minimized these distances, as the path along the density crest is the one most centered in relation to all cells.

Our method takes as input the following parameters:

1) The approximate locations of the starting and ending trajectory points. The method is robust to some deviation from the true starting and ending cells, since clusters of similar cells are segmented together anyway, and even trajectory paths originating from relatively distant starting points converge at some point along the density crest.

2) The threshold above which the difference in neighboring densities justifies a detour from the direction aligned with the moment. A lower threshold is preferable for cases where the marker landscape has high curvature (e.g. circular), and sticking with moment alignment might significantly divert the trajectory form passing along the density crest. However, a lower threshold is prone to trajectory off-track, as small fluctuations in densities can lead to superfluous coiling. It is thus wise to consider some degree of smoothing during pre-processing, which also helps remove redundant anchors. However, in our case no smoothing was necessary as part of data pre-processing, as superfluous anchors were removed successfully by the method described above, and no significant trajectory off-tracking was observed.

3) The grid density. A denser grid allows a compact trajectory, resulting in a higher resolution. Computational expense of a dense grid becomes pronounced as more dimensions are added, since the number of grid points rises exponentially with the number of markers ($\sim Grid\ density^{(\#\ of\ markers)}$). We therefor chose a denser grid only when necessary, and used only the 8 most valuable markers in each segment for sub-segmentation (Section 7).

4) The vicinity of neighbor search during calculation of the next step. When deciding on the position of the next trajectory points we only looked at the closest nearest neighbors. However, a more complex version could include searching for neighbors at further circles (preferably aligned with the moment), and allowing more flexibility during progression within this larger circle towards the chosen neighbor. See Section 7 below for parameter choice for the results in the main text.

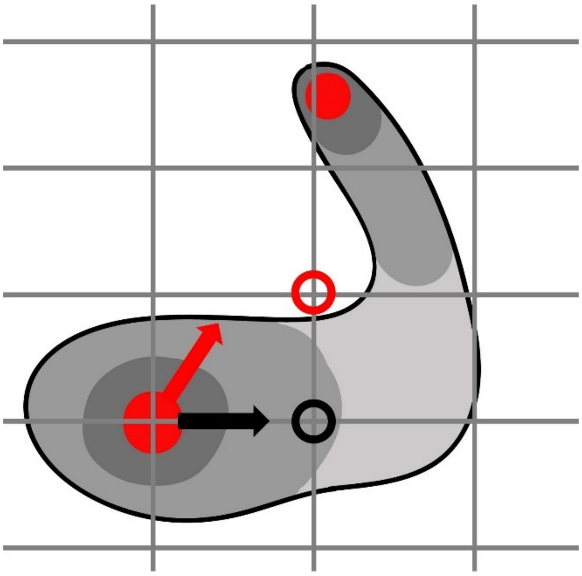

Figure S2: Deviation from progression towards moment direction. Sketch shows some imaginary density topology with two local maxima in the red regions. Gray shades representing lower cell density (lowest density is in lighter gray or white regions). In this cartoon, the moment is roughly aligned with the red arrow (pointing at the next maxima). Progression in the moment direction would lead to a grid point positioned in a very low density region (encircled in red), whereas allowing a detour in the direction of the black arrow allows a better overlap of the trajectory with the density crest. In this example, the density difference between grid points encircled in red and black might exceed a predetermined value and result in a better trajectory fit.

### Section 3: Time calculation

To derive the equation for time let us first define the following notations:

$l$ – The trajectory index, starting at $l = 1$ and ending at some $l = l^{final}$. Each index corresponds to a point in high dimensional space, and effectively reduces dimensionality to 1. Every cell is affiliated to the closest point on the trajectory based on Euclidian proximity.

$f_i$ – Cell density in the $i^{th}$ segment: $\frac{Number\ of\ cells\ closest\ to\ i^{th}\ index\ of\ l}{Total\ number\ of\ cells}$ (has units of $[\frac{1}{l}]$).

$F_i$ – The cumulative density up to the $i^{th}$ segment: $F_i = \int_{l=1}^{i}(f_l)dl$.

$v_i$ – The progression speed along the $i^{th}$ segment defined as $\frac{dl^i}{dt}$.

$\gamma_1^i$ – The proportion of cells dividing in the $i^{th}$ segment per time unit (has units of $[\frac{1}{t}]$).

$\gamma_2^i$ – The proportion of cells undergoing apoptosis in the $i^{th}$ segment per time unit (has units of $[\frac{1}{t}]$).

$T$ – The cycle length (time at $l = l^{final}$), which we assume is known.

The continuity equation in its general form is given by:

$$\frac{df}{dt} + \nabla \cdot J = \sigma$$

Where $f$ is cell density, $\nabla$ is the divergence operator, $J$ is the density flux, and $\sigma$ is cell-density generation per time unit. At steady state, $\frac{df}{dt} = 0$, as local cell density is constant in space. Using the notations above, density flux can be written as

$J = f \cdot v$. The sources for generation and degradation during the process are division and apoptosis, hence $\sigma = (\gamma_1 - \gamma_2) \cdot f$. Thus, at the level of each segment, using index form, continuity equation implies that $\nabla(f \cdot v) = (\gamma_1 - \gamma_2) \cdot f$ (Figure 2B).

There are three cases worth considering when solving this equation:

_No division or apoptosis_

In this case $\gamma_1 = \gamma_2 = 0$ (or more generally $\gamma_1 = \gamma_2$) , and the equation can be simplified to $\nabla(f \cdot v) = 0$.

$$f \cdot v = f \cdot \frac{dl}{dt} = const$$

$$t = const_1 \cdot \int fdl = const_1 \cdot F + const_2$$

Since at $F = 0$ also $t = 0$, and at $F = 1$ by definition $t = T$, the private case of no division or apoptosis can be compactly solved as:

$t(i) = F(i) \cdot T$, where $i$ is the trajectory index.

*Constant division and apoptosis*

In this case $\gamma_1 - \gamma_2 = constant$ throughout the process ($constant$ >0). Now:

$$f \cdot v = (\gamma_1 - \gamma_2) \cdot F + c1$$

$$t = \int \frac{f}{(\gamma_1 - \gamma_2) \cdot * F + c1} dl = \frac{1}{(\gamma_1 - \gamma_2)} \int \frac{f}{F + c2} dl =$$

$$= \frac{1}{(\gamma_1 - \gamma_2)} \ln(F + c2) + c3 = \frac{1}{(\gamma_1 - \gamma_2)} \ln\left(\frac{F + c2}{c4}\right)$$

Where $c1, c2, c3, c4$ are all constants. Appling the same boundary conditions as before at $F = 0$ and $F = 1$, we get:

$$t = \frac{1}{(\gamma_1 - \gamma_2)} \ln\left(\frac{F + c}{c}\right), \;\; c = \frac{1}{e^{(\gamma_1 - \gamma_2)T} - 1}$$

*Varying division and apoptosis rates*

In this case, $\gamma_1$ and $\gamma_2$ have different values in each segment, necessitating a numeric approach:

At the level of the first segment, continuity implies that $f_1 \cdot v_1 - \varphi = (\gamma_1^1 - \gamma_2^1) \cdot f_1$ (upper indices represent trajectory index), where $\varphi$ is the flux of entering cells.

At the level of any other segment, continuity implies that:

$f_l \cdot v_l - f_{l-1} \cdot v_{l-1} = (\gamma_1^l - \gamma_2^l) \cdot f_l$, or:

$$f_l \cdot v_l = \varphi + \sum_{i=1}^{l} (\gamma_1^i - \gamma_2^i) \cdot f_i$$

Since $v_l = \frac{dl}{dt(l)}$, the time at each trajectory location is:

$$t(l) = \sum_{i=1}^{l} \frac{f_i}{\varphi + \sum_{j=1}^{i} (\gamma_1^j - \gamma_2^j) \cdot f_j}$$

$\varphi$ can either be determined experimentally, or calculated numerically using the estimated time at a given point on $l$ (e.g. estimated $T = t(l)$).

Notably, the only source of cell removal accounted for during the process is apoptosis. When the process ends, that is when each cell reaches its $t = T$, cells leave the thymus and enter the circulation. Towards the end of the trajectory, some cells early to mature leave the thymus while other cells require more time to do so (as implemented in our model by adding noise to $T$). Hence, at these trajectory regions, local cell density is thinned by cells exiting the thymus and not via apoptosis contribution alone. Since we do not 'correct' $\gamma_2$ for this other contributory source of cell removal, some deviation can be noted towards the end of the trajectory when comparing average simulated time (the true time) to calculated time using the continuity principle (Figure 2C).

### Section 4: Division and apoptosis rates assessment

Plotting $\log(IDU)$ as a function of $DNA$ expression reveals two distinct clusters of cells, with low and high $IDU$ levels, which allows easy gating of dividing cells within the time that passed between $IDU$ injection and mouse harvesting ($\delta t$) (Figure 3B in the main text). To estimate the proliferation parameter in each segment ($\gamma_1$), we divide the proportion of gated $IDU$-positive cells in the segment by $\delta t$.

$Caspase3$ positive cells could also be distinguished as the upper separate layer of cells expressing higher levels of the marker (Figure 3C in the main text and Figure S3 panels A,C). Estimating the proportion of cells that divide per time unit in each segment ($\gamma_2$) is more complicated, since the time framework in which apoptosis occurs is less implicit. To this end, we only gated cells that were positive to both $Caspase3$ and $IDU$, allowing adoption of the same time interval $\delta t$ as when calculating $\gamma_1$. Since gating cells positive to both markers could potentially lead to selection bias, underestimating the apoptosis rate in each segment, we checked how $Caspase3$ positive cells distributed in the two $IDU$ groups ($IDU$-high and $IDU$-low). A skewed distribution of $Caspase3$ positive cells, e.g. towards the $IDU$-low cluster, would indicate underestimation of cells in the apoptosis process when gating cells positive to both markers. Luckily, $Caspase3$ distributed roughly evenly (Figure S3 panels B,D), indicating that the gated cells positive to both markers are suitable for assessment of the parameter $\gamma_2$.

As described in the main text, about 90% of T-cells are known to be removed from the thymus through 'death by neglect', which is undetected using standard apoptosis markers. We hence increased $\gamma^2$ in a couple of segments (indicated by the arrow in Figure S3 panel E), so that 90% of the cells were lost. Direct measurement of cell numbers in the adoptive-transfer experiment (Figure 5A) verified a reduction of ~90% in T-cell numbers around day 21 post injection (Figure S3 panel F).

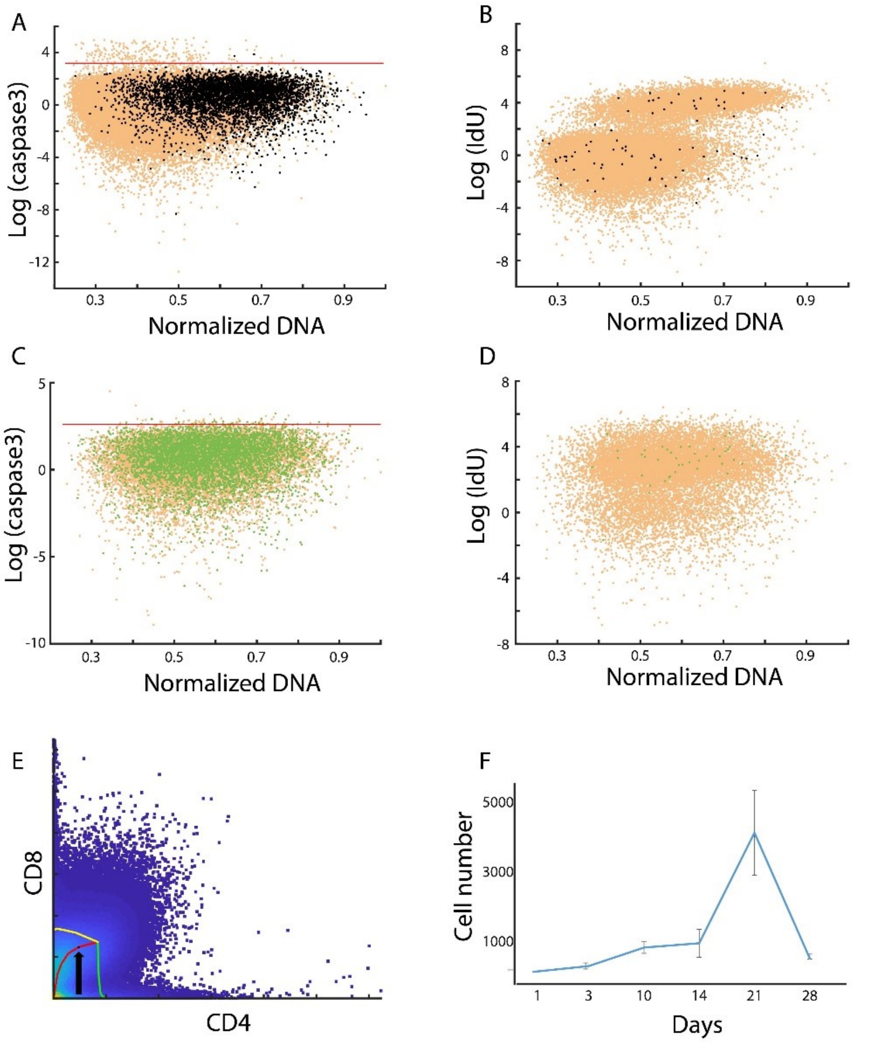


Figure S3: Assessing proliferation and apoptosis rates. (A,B) Harvesting 1-hour post IdU injection. Cells colored black in (A) are IdU positive gated cells (~4.5% of all cells). Cells above horizontal red line are caspase3 positive. IdU positive cells that were also caspase3 positive were considered as cells that will undergo apoptosis within 1 hour. Black cells in (B) are caspase3 positive cells, which are roughly evenly distributed between clusters of IdU positive cells (0.33 % of cells in upper cluster) and IdU negative cells (0.25% of cells in lower cluster). (C,D) Harvesting 6-hours post IdU injection. Here green cells represent IdU positive cells in (C) (~24.2 % of all cells) and caspase3 positive cells in (D). Notably proportion of IdU positive cells 6-hours post injection is approximately six times compared to 1-hour post injection, resulting robust proliferation rates after normalizing by $\delta t$. (E) Arrow pointing at the position along the trajectory of one of the mice where $\gamma^2$ was increased so that 90% of the cells were removed. (F) Cell numbers 1,3,10,14,21 and 28 days post adoptive-transfer. A significant drop in ~90% in cell number is detected around day 21 at the end of the DP stage where 'death by neglect' is known to occur.

### Section 5: A two-step segmentation process

Segmentation involves assigning each cell to a trajectory point that is in closest Euclidian vicinity. Assigning cells using multiple markers while giving each marker the same weight, can result in a dispersed affiliation pattern when projecting on CD4/CD8 space (Figure S4A).

Dispersion in CD4/CD8 space can result e.g. in instances where cells in DN phase are assigned to trajectory points located in the DP region, and should therefore be avoided. To solve this, we conducted segmentation in two steps, where at first we performed initial cell-assignment based on CD4 and CD8 alone (Figure S4B). In the second step, we further performed sub-segmentation in each segment, this time adding more markers (Figure 4A,B in the main text). The markers chosen for expanding each segment were those with highest local variability (aside from CD4 and CD8 which were always accounted for). Hence, dynamics could be unfolded without the expense of scattering in CD4/CD8 space.

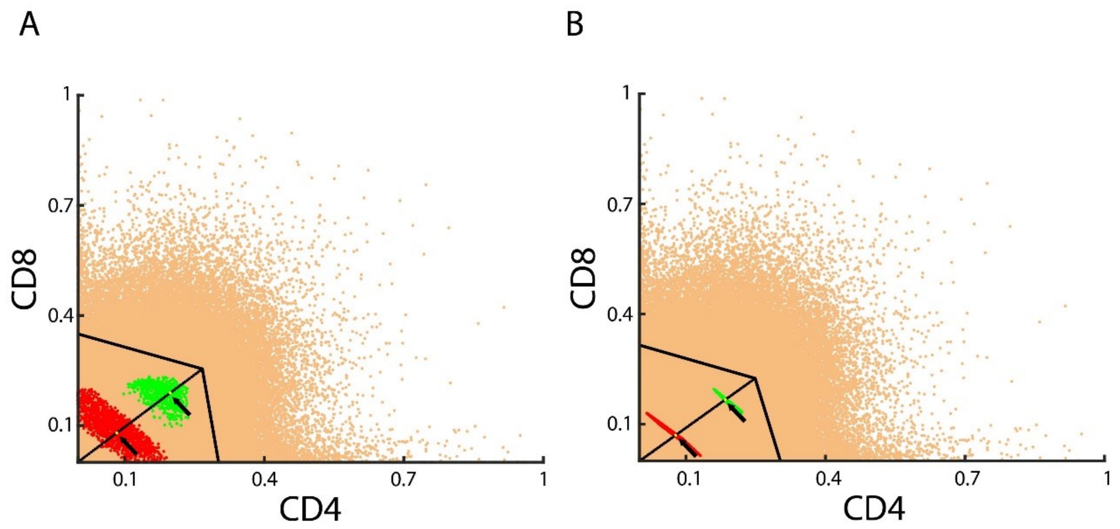


Figure S4: (A) Cell dispersion. Cells were assigned to each trajectory point using the 8 following markers: CD4, CD8, CD45, CD24, CD25, CD3, CD117, CD44. Figure shows projection upon CD4/CD8 space, where cells colored in red and green are those closest to trajectory points highlighted by arrows. (B) Assigning cells in 2 dimensions results in a compact affiliation pattern, avoiding dispersion when later adding more dimensions.

### Section 6: Profiles of all markers

In this section, we compare marker expression profiles along the trajectories leading from DN (specifically DN3) and DP to CD4 SP and to CD8 SP. The trajectories overlap up to the bifurcation point.

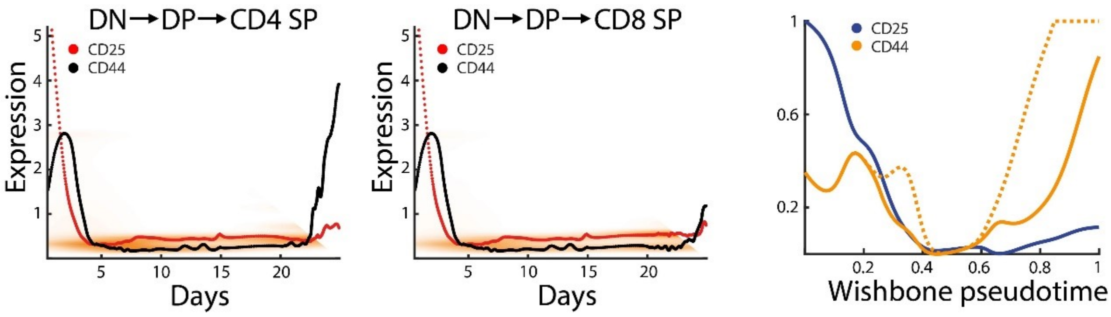


Figure S5: DN markers along the trajectory leading to CD4 SP and CD8 SP (left and middle panels). Right panel presents Wishbone pseudotemporal trajectory for these markers.

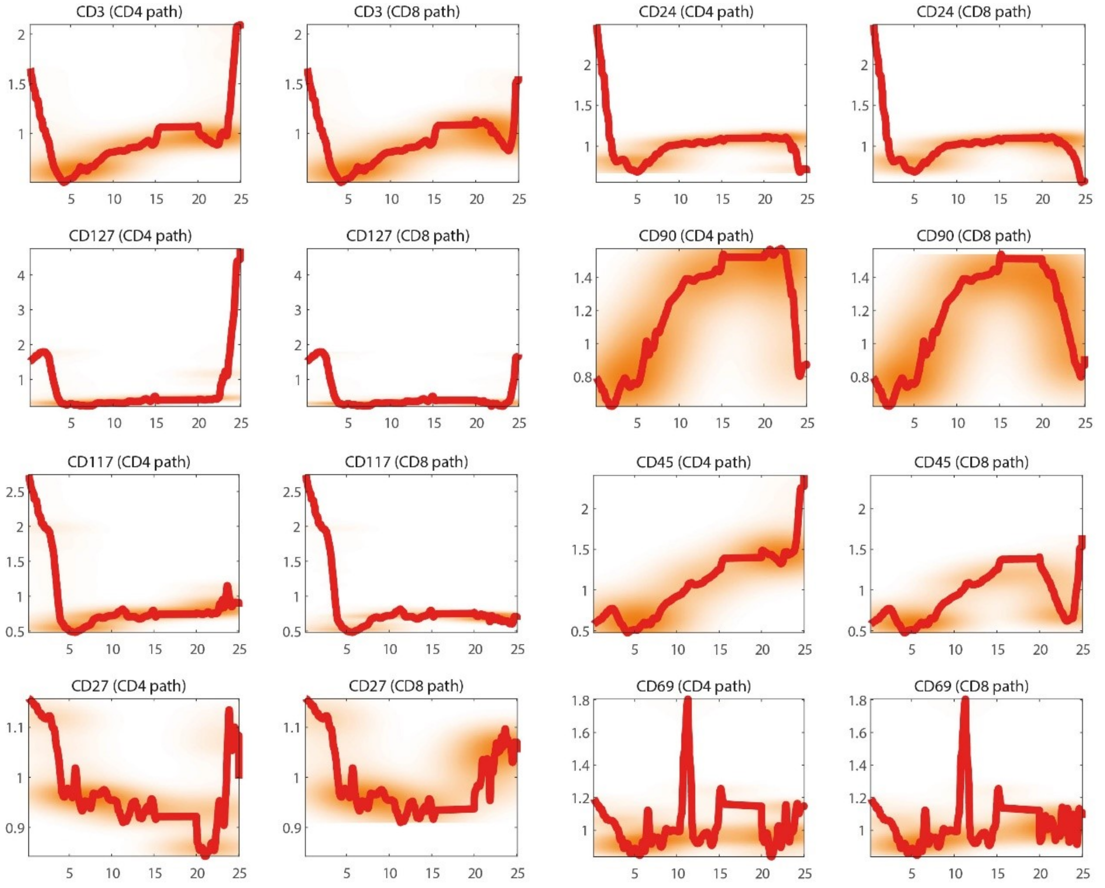

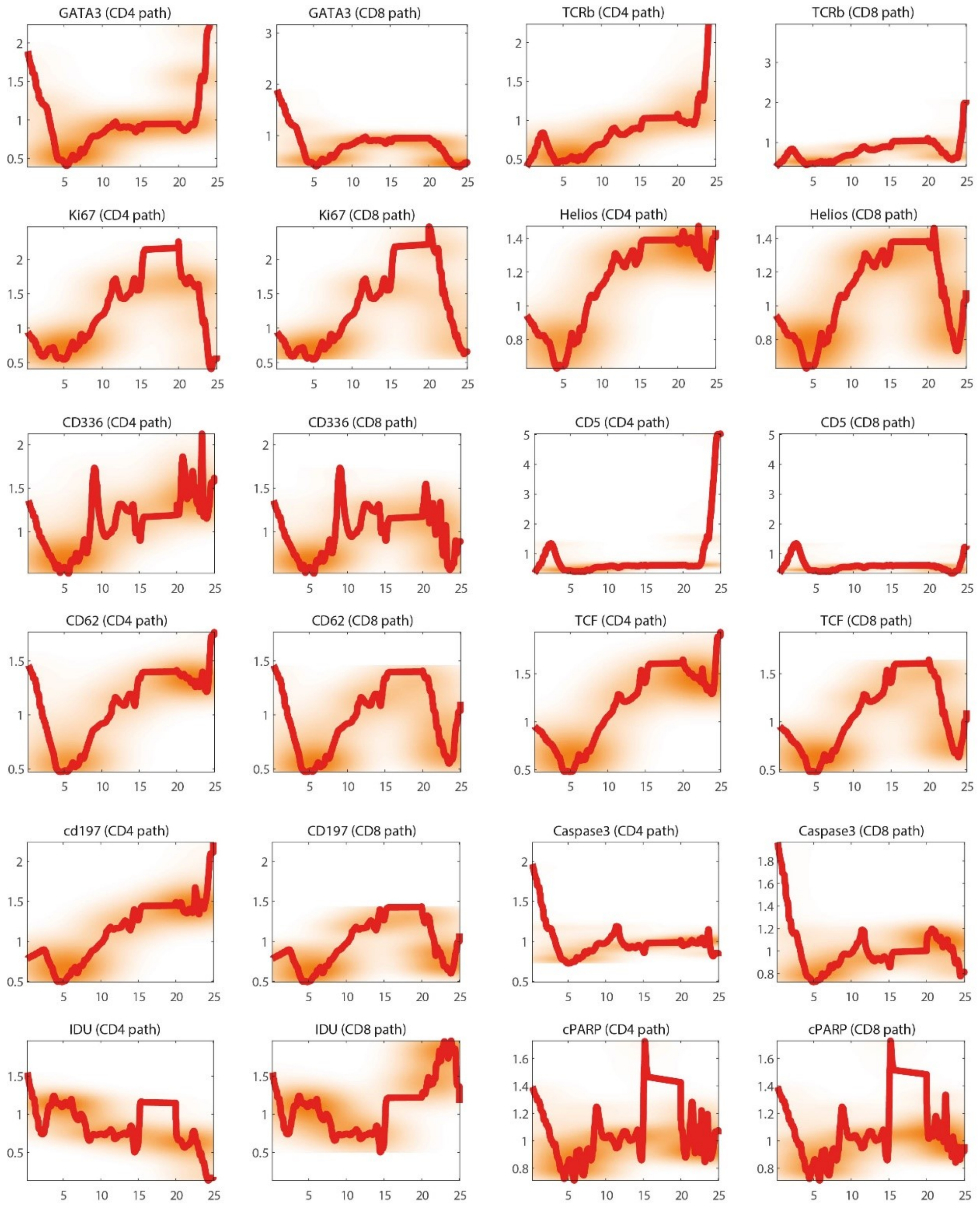
GATA3 (CD4 path)
GATA3 (CD8 path)
TCRb (CD4 path)
TCRb (CD8 path)
Ki67 (CD4 path)
Ki67 (CD8 path)
Helios (CD4 path)
Helios (CD8 path)
CD336 (CD4 path)
CD336 (CD8 path)
CD5 (CD4 path)
CD5 (CD8 path)
CD62 (CD4 path)
CD62 (CD8 path)
TCF (CD4 path)
TCF (CD8 path)
cd197 (CD4 path)
CD197 (CD8 path)
Caspase3 (CD4 path)
Caspase3 (CD8 path)
IDU (CD4 path)
IDU (CD8 path)
cPARP (CD4 path)
cPARP (CD8 path)

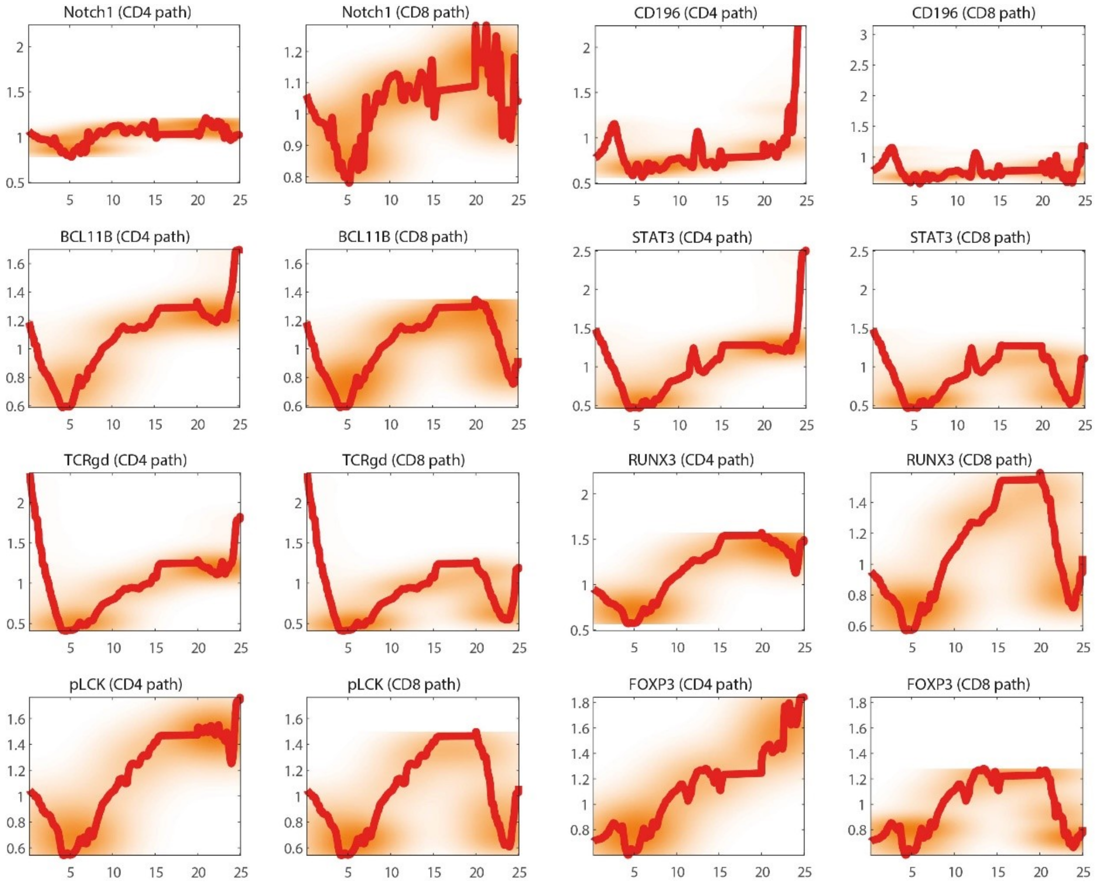


Figure S6: Different markers plotted as function of real time. Vertical axis is the expression and horizontal axis is time (days). Marker values are plotted twice, along the CD4 and CD8 paths (up to the bifurcation point the plots overlap).

### Section 7: Data processing and parameters

Here we first describe the pipeline for pre-processing data before inferring the trajectory path and calculating time. We later give the parameters we chose for results presented in the main text and sections above.

**Data pre-processing**

Pre-processing the data before trajectory inference included the following steps:

1) **Raw data:** Raw data from a single mouse thymus, including expression levels of unsynchronized differentiating single-cells measured by CyTOF, was loaded to a custom written Matlab program. We analyzed altogether 12 mice, including mice harvested 1, 4, 6 and 12 hours after IdU injection (3 mice in each group). Initial cell counts for each mouse ranged between 40,000 to 174,000 (with a mean of 97,000 and a standard error of 16,500). Notably, cells were not synchronized, representing all stages of maturation in the thymus.

Expression levels of the following markers were measured for every cell:

| | | | | | | | | |
|---|---|---|---|---|---|---|---|---|
| CD4 | CD8 | IdU | Caspase3 | CD45 | CD24 | cPARP | IKAROS | CD69 |
| Notch1 | TCF7 | CD90 | CD25 | CD3 | CD336 | BCL11B | CD62L | pLCK |
| FOXP3 | GATA3 | CD5 | Ki67 | RUNX3 | CD90 | TCRgd | CD117 | STAT3 |
| CD27 | TCRb | CD69 | CD44 | TCF7 | CD197 | CD127 | Helios | CD196 |
| DNA1 | DNA2 | Cisplatin | NK1 | CD11b, CD11c, CD19 (Non T-cells markers) | | | | |

Table 1: Markers measured by CyTOF for each cell.

2) **Gating:** To gate out only T-cells from the raw data, we first used the inverse hyperbolic sine transformation, $Marker \rightarrow arsinh(\frac{Marker}{cofactor})$, choosing $cofactor = 5$. T-cells were gated by choosing cells with low levels of 'Non T-Cells' markers (see Table 1), with the transformed marker threshold set at ~5 (Figure S6).

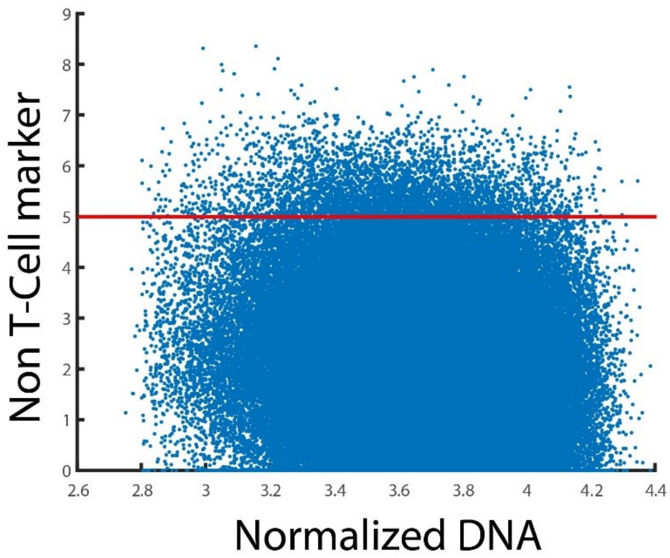


Figure S7: Gating T-cells. T-cells were chosen if levels of the 'Non T-cell' marker was lower than the threshold (horizontal red line at value ~5 depending on mouse). 'Non

T-cell' marker levels were first transformed via the inverse hyperbolic sine function as described above.

3) **<u>Outlier removal</u>:** we removed any cell with expression levels above the 99.99 percentile, for any of these markers:

CD4, CD8, CD45, CD27, CD69, CD90, CD24, CD25, CD3, CD336, CD62l, CD5, CD117, CD44, CD197, CD127

4) **<u>Normalization</u>:** Normalizing each marker by its mean value was performed, since marker value outputs from CyTOF are arbitrary and are only significant for comparing signaling strengths between cells for a given marker, but not for comparing different marker values.

5) **<u>Parameters:</u>**

The trajectory was initially calculated in two-dimensions (CD4/CD8 space) as described, where the trajectory was divided into 60 evenly-spaced points from the initial starting point until the point of bifurcation. Using the same interval between points, the trajectory was extended from the bifurcation point along each of the SP states. A detour from the path leading to the next anchor was allowed if the density in a neighboring position on the grid was 1.5 times that of the mean of all neighboring cells and twice that of the density in the default point.

After the trajectory position was determined, cells were assigned to the closest trajectory point based on Euclidian distance, and the time equation was solved numerically (Figure 2B and SI section 3). For sub-segmentation we picked the 8 most varying markers in each segment. We chose a different grid density for each segment based on the time spent in the segment that was calculated in two dimensions (Figure 4). The maximal allowed grid size in 8-dimensions was of size 10 for the segment where the maximal time was spent, and other segments were assigned grid sizes proportional to the ratio in times compared to the maximal time.
